# Enhanced conspicuousness of prey in warmer water mitigates the visual constraint of turbidity for predators

**DOI:** 10.1101/2024.09.17.613492

**Authors:** Costanza Zanghi, Jolyon Troscianko, Christos C Ioannou

## Abstract

1. Changes in environmental conditions impact predator-prey interactions by altering behaviour through sensory and non-sensory pathways. Elevated water temperature and turbidity are known to alter activity levels and anti-predator responses in prey fish, and are increasing globally as a result of anthropogenic activities. Less is known about how temperature and turbidity impact predators’ ability to detect prey directly, or indirectly via changes to prey behaviour.
2. We quantified the detectability of Trinidadian guppies (*Poecilia reticulata*) free-swimming in a large arena from the perspective of a stationary visual predator (simulated as an underwater camera). We used a fully factorial experimental design testing the independent and combined effects of increased temperature and turbidity.
3. We found that both stressors had a strong influence on the appearance of prey (objectively quantified as the mean magnitude of the optical flow in the videos). As expected, turbidity reduced the frequency of detection between the guppies and the simulated predator, i.e. the magnitude of optical flow exceeded the threshold for a ‘detection event’ more often in clear water. Events were also shorter in duration in turbid water, reducing the time available for a predator to detect the prey. However, during an event, prey were more detectable in warmer water (i.e. the mean magnitude was greater).
4. Although we found no evidence of interactive effects of turbidity and temperature on the response variables, their cumulative main effects suggest an antagonistic effect on the overall rate of prey capture.

## Introduction

An ambush predator must first encounter, detect, and recognise potential prey items before launching an attack (Lima & Dill, 1990). Encounters are determined by the rate at which predator and/or prey are within the sensory range of the other. Studying encounter rates thus often involves exploring the factors that allow a predator to get within physical proximity to their prey, and conversely, the avoidance behaviours of prey that minimise encountering predators (Ferrero et al., 2011; Ioannou et al., 2008). Detection is the sensory process that happens within an encounter, resulting in the identification and localisation of an object (the signal) that is not part of its surrounds (the noise) (Troscianko et al., 2009). Once detected, the object must be classified as something of interest, which can either result in successful recognition of the object, or misidentification (as occurs with mimicry or masquerade, Ruxton et al., 2018). Each of the steps in the predator-prey interaction sequence (Lima and Dill, 1990) can be influenced by various factors including the hunting mode of predators (Sims et al., 2008; Wearmouth et al., 2014), the prey’s foraging and anti-predator strategies (Carroll & Sherratt, 2013; Magurran, 1990; Turesson et al., 2009), other predator and prey traits (Franco et al., 2022; Harris et al., 2010; Szopa-Comley et al., 2020) and abiotic environmental conditions (Abrahams et al., 2007; Domenici et al., 2019).

Environmental conditions can impact encounter rates by altering how predators and prey use and move through their environment. For example, turbidity can alter the search behaviour of diving seabirds that use vision to pursue prey underwater (Darby et al., 2022). Similarly, bat species whose prey-detecting echolocation calls overlap in frequency with sources of anthropogenic acoustic noise reduce activity around such sources (Bunkley et al., 2015). When predator and prey become close enough that at least one can detect the other, i.e. an encounter occurs, environmental conditions can affect the probability of detection by masking sensory cues. For instance, visual predators such as the three-spined stickleback (*Gasterosteus aculeatus*) were less likely to respond to prey in environments with increased visual noise (i.e. caustics; Attwell et al., 2021). Similarly, environmental conditions can distract individuals, impairing their ability to assess their surroundings efficiently. This distraction can weaken anti-predator responses, as seen in hermit crabs (*Coenobita clypeatus*) when exposed to both acoustic and visual noise (Chan et al., 2010), or in dwarf mongooses (*Helogale parvula*), which were slower to respond to secondary predator cues under anthropogenic acoustic noise (Morris-Drake et al., 2016). Increased turbidity can impact predator-prey interactions by both masking visual cues, reducing detection distances (Nieman & Gray, 2019) and disrupting shoal dynamics (Borner et al., 2015), and diminishing prey’s risk awareness, thereby distracting them from threats (Gregory, 1993). Additionally, environmental parameters that affect the physiology and behaviour of prey can render them more or less conspicuous to their predators. Predators detect prey more easily when their activity levels are raised due to higher water temperatures (Krause & Godin, 1995), because predators have often evolved to be highly sensitive to prey movement (Ioannou & Krause, 2009), and movement is more readily detectable than other visual information (Troscianko et al., 2009). This explains why, for instance, after being detected by a predator, freezing is a common response observed in many taxa (Eilam, 2005).

Little is known about how the interactions between multiple environmental stressors affect encounter and detection in predator-prey interactions. This is important because rapid environmental change is affecting predator-prey interactions at a global scale (Abrahams et al., 2007; Domenici et al., 2007). Environmental stressors, here defined as any parameter experienced by a system with increased frequency and intensity (Townsend et al., 2008), rarely impact biological processes in isolation. Instead, they co-occur, leading to cumulative impacts on a system (i.e. additive effects). Additionally, these stressors can interact in ways that cause “ecological surprises” (Paine et al., 1998). In such cases, the resulting impact is unpredictable based on the responses to each stressor in isolation, resulting in an overall response that is greater (synergism) or lesser (antagonism) than the sum of the individual responses (Folt et al., 1999).

In this study, we adopted an approach developed by Turesson and Brönmark (2007), who assessed encounter rates between prey fish and a side-view facing underwater video camera, which was used to quantify prey appearance from the perspective of an aquatic ambush predator. Based on their observations in natural lakes, Turesson and Brönmark (2007) found that eutrophication and water transparency had important effects on encounter rates. In more turbid conditions, the distance at which prey were visible was so impaired that the probability of encountering prey decreased, even when prey density was at its highest. While Turesson and Brönmark (2007) directly observed video recordings to extract behavioural metrics such as camera detection distance, the frequency of encounter, encounter duration and shoal size, here we used optical flow analysis to objectively estimate the apparent motion of prey fish from the video footage as a measure of the prey’s likelihood of being encountered and then detected by a predator. Optical flow analysis is a computer-vision tool widely used for detecting and quantifying motion (Sun et al., 2014). Its application in biology spans diverse fields, including the assessment of farmed animal welfare (Dawkins et al., 2009, 2012), the investigation of animals’ self-motion and spatial awareness (Alexander et al., 2022), and the analysis of footage from wild animal camera traps (Khorrami et al., 2012). Optical flow allowed us to quantify not only the frequency and duration that prey could be detected (as in Turesson and Brönmark (2007)), but also how conspicuous prey are during these encounters. All of these factors can be considered to contribute to the detectability of prey, and be likely indicative of the frequency of attacks by predators and hence wider ecological impacts of predation (Chase et al., 2002; Lima, 1998).

To investigate the effects of changing environmental conditions on prey detectability, we designed a fully factorial laboratory experiment. We used free-swimming Trinidadian guppies (*Poecilia reticulata*) under control and elevated water temperatures and turbidities, as perceived by a hypothetical stationary predator. Increased turbidity reduces light levels and visual contrast, limiting the detection abilities of predators (Beauchamp et al., 1999; Hansen et al., 2013; Vogel & Beauchamp, 1999). In addition to the direct effect of turbidity on visibility underwater, these stressors can have contrasting effects on the activity and shoaling behaviour of guppies, which, in turn, impact encounter rates. The shoaling behaviour of guppies in warmer temperatures have been observed to vary, increasing or decreasing, depending on the level of perceived risk (Weetman et al., 1998, 1999). Likewise, other studies have reported varying effects of turbid water on guppies’ activity levels, with some finding no effect (Zanghi et al., 2023), while others have observed heightened or lessened activity (Borner et al., 2015; Ehlman et al., 2015).

Building on this previous literature, we hypothesised that in warmer water, guppies would be more active and hence more frequently observed within the camera’s field of view. This would, however, reduce the amount of time over which detection could occur for each event (limiting the probability of detection per event). Under turbid conditions, detection distances are reduced, but moving targets closer to the camera within its field of view are still detectable. Thus, while encounter frequency would be reduced, detection probability at closer distances would increase. In the interaction treatment, we hypothesised a synergistic effect, where guppies entering the camera’s field of view would be more detectable compared to the additive effect of turbid and warmer water in isolation. However, if temperature and turbidity have opposing effects on activity, in the interaction treatment there may be an antagonistic effect, where the detectability is similar to the control treatment of clear and ambient temperature water.

## Methods

### Experimental set up

The data presented in this study was collected during a parallel study by Allibhai et al. (2023), where details on the provenance and housing of the study subjects are described. In brief, 720 adult, mixed-sex guppies (*Poecilia reticulata*) were haphazardly assigned to four 200 L holding tanks. Over 16 days between February and March 2021, all guppies were tested once under each of four experimental treatments: control (same as holding conditions: 0 Nephelometric Turbidity Units, NTU and 22⁰C), increased turbidity (5 NTU, 22⁰C), warmer temperature (0 NTU, 29⁰C), and a combined turbid and warmer treatment (5 NTU, 29⁰C). These conditions reflect the natural variation observed in the guppies’ native habitat in Trinidad (Zanghi et al., 2024). The sequence of treatments was arranged in a balanced crossover Latin square design to reduce order effects (Figure 1a; N_trial_=91; five trials (two warm, two turbid, and one in the interaction treatment) were excluded as the camera malfunctioned during filming).

**Figure 1.**
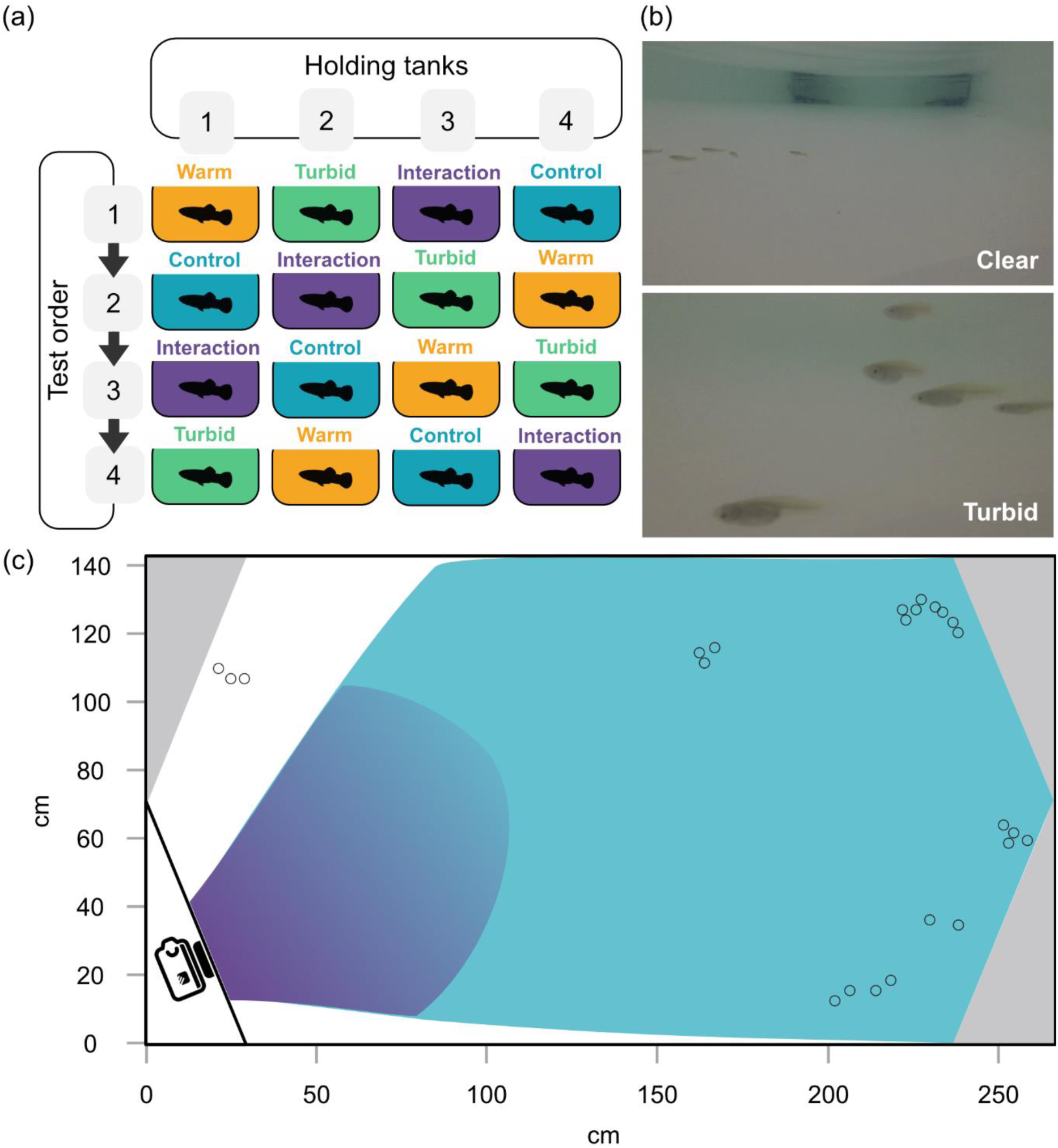
Protocol detailing the balanced crossover Latin square design for treatment order **(a)**: control (0NTU, 22⁰C), turbid (5NTU, 22⁰C), warm (0NTU, 29⁰C), and interaction (5NTU, 29⁰C). **(b)** Camera views in clear and turbid water. **(c)** Schematic of the experimental arena showing refuges housing the heaters (grey corners), example of fish positions in a single frame (open circles), and the transparent divider (black line) for the camera placement. The lighter blue area represents the field of view of the camera in clear water, the darker blue in turbid water. Adapted from Allibhai et al., 2023.

For each trial, approximately 30 fish were haphazardly caught with hand nets from the holding tank and gently released in the middle of the experimental arena (Figure 1c, 255 x 140 cm, with 10 cm water depth). After a 4-minute acclimatisation period, fish were filmed from the side view for 11 minutes using a camera underwater (GoPro Hero5) placed behind a transparent divider in one corner of the tank. Trials were recorded in 2.7K resolution (2704 x 1520 pixels), in wide mode, and at a frame rate of 30 frames per second.

Turbidity was achieved using dissolved kaolin clay powder (Figure 1b) and it was measured using a turbidity meter (Thermo Scientific Orion AQUAfast AQ3010) before and after each trial with increased turbidity. Temperature was manipulated by using submersible heaters placed in the corner of the experimental arena, and it was measured using a thermometer before and after each trial.

### Data extraction

Each video was processed using Matlab R2021a (The MathWorks Inc., 2021). Videos were trimmed at 19800 frames (11 minutes) from the last frame to ensure an equal sample size among trials. Each frame was then converted to greyscale and optical flow was calculated using the estimatedFlow() function based on the Lucas-Kanade method (Barron et al., 1992). This method calculates the magnitude of the pixels’ flow vector from one video frame to the next. The flow vector consists of the horizontal and vertical components, which indicate displacement of a pixel in the X and Y directions respectively, while the magnitude of the flow vector represents the intensity of the flow at that pixel. For each video frame, magnitude was averaged (mean) across all pixels, yielding one measure of magnitude per frame (N_Frame_ across all trials = 1,801,800) (Figure 2). While these algorithms do not directly replicate predator motion vision, their ability to detect contrasts in moving prey should correlate with real visual systems, albeit with different detection thresholds that are yet to be well-characterised for guppy predators.

**Figure 2.**
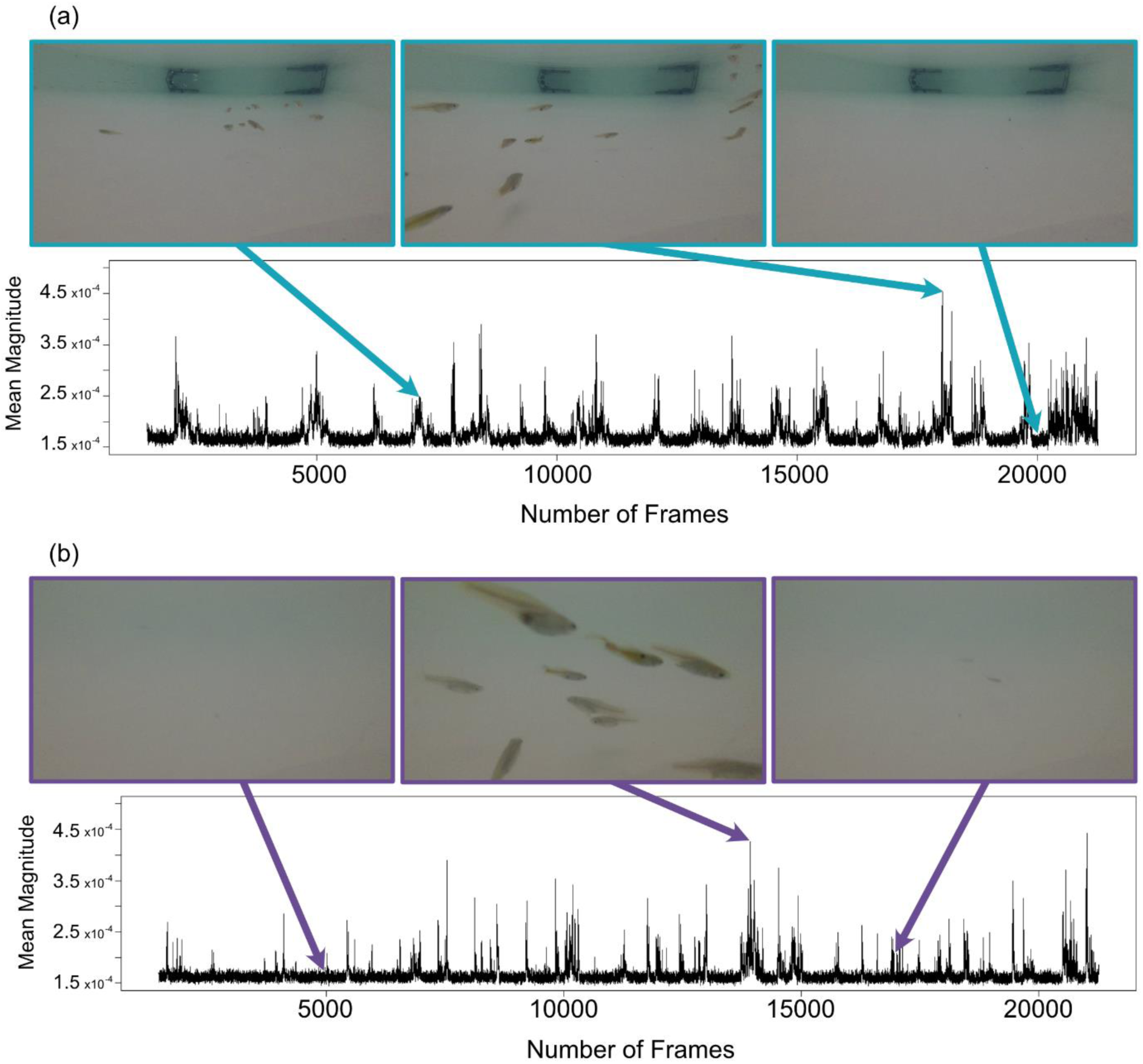
Example from two trials, for the control (**a**) and interaction (**b**) treatment. In the time series, the dark band at low mean magnitudes represents background noise, while spikes in mean magnitude represent frames in which fish motion was detected. Three specific frames and their respective mean magnitudes are highlighted within each trial.

Frames where the mean magnitude exceeded a threshold were considered occurrences where the conspicuousness of the fish (i.e. detectability based on the optical flow) was great enough to be detected by a predator. The statistical analysis was carried out on different datasets which varied in the threshold that was applied, and the results compared qualitatively. The thresholds were calculated as the 99^th^, 95^th^, 90^th^, 80^th^ and 70^th^ percentiles of the mean magnitude pooled across all trials (Figure 3). We followed this approach to test how general the results were across different thresholds as a sensitivity analysis. Additionally, although the relationship between optical flow as measured in our experiment and detectability to a real predator of the guppy is unknown, varying the threshold could be analogous to comparing predators with varying visual sensitivities. Here, the highest threshold is comparable to predators with lower sensitivity and only able to detect high levels of motion, while the lower thresholds represent predators able to detect subtle prey motion.

**Figure 3.**
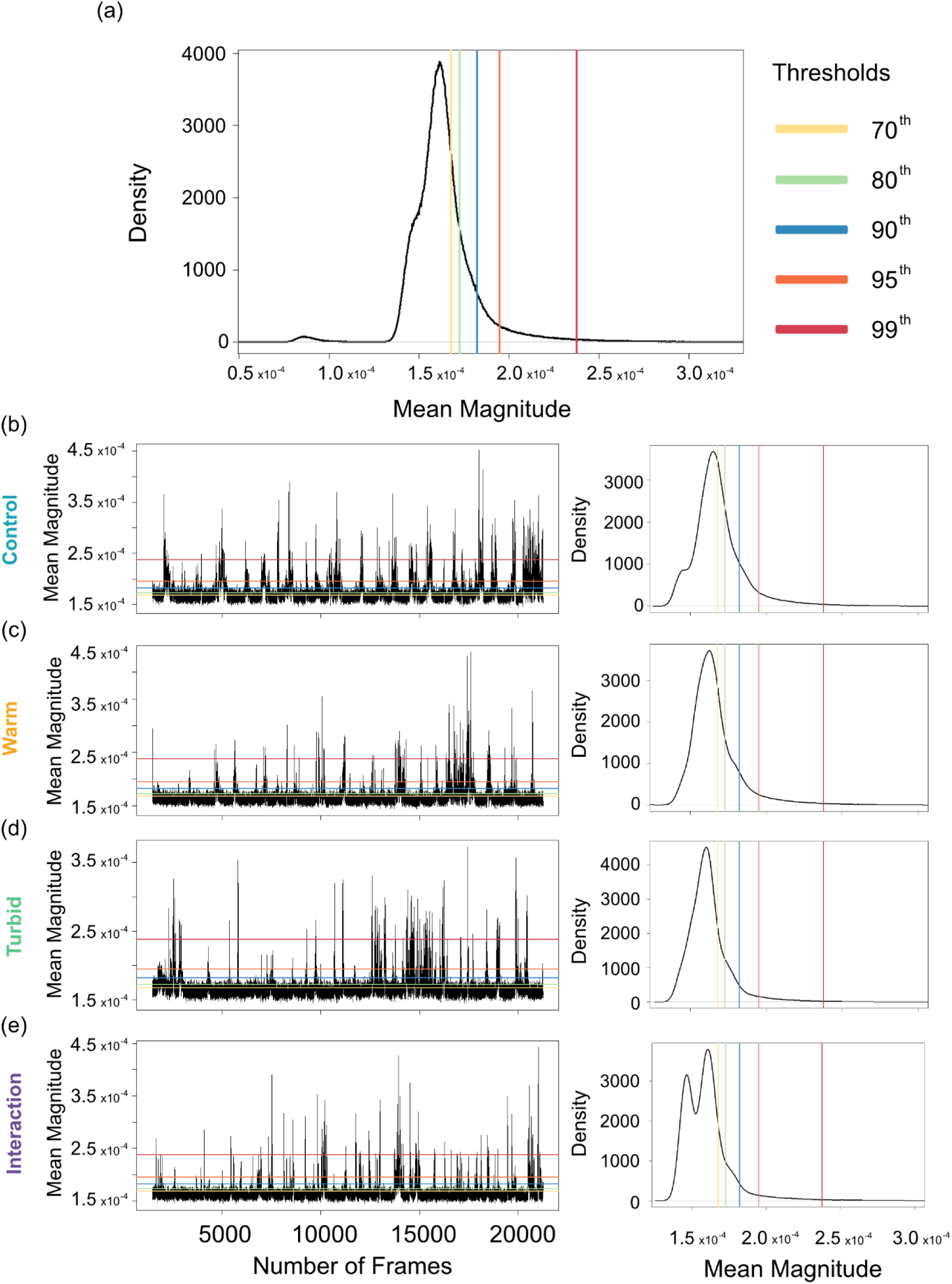
Thresholds of mean magnitude for the aggregated data; N=1,801,800 (**a**). Examples of mean magnitude across one individual trial for the control (**b**), warm (**c**), turbid (**d**) and interaction (**e**) treatment, alongside the mean magnitude distribution for each treatment (N_cont_=470,249; N_warm_=430,651; N_turb_=450,450; N_int_=450,450).

For each trial we calculated the proportion of frames where the mean magnitude exceeded the threshold, where a higher proportion indicates trials where fish were detectable more often within the field of view of the camera. The mean magnitude recorded for each frame above the threshold was averaged (mean) per trial, where a greater mean magnitude indicates prey that were more detectable.

Consecutive frames where the mean magnitude exceeded the threshold were grouped as a single “detection event”, which corresponds to the period of time from fish swimming into the visual field of the camera to when they were no longer visible. The number of detection events was then calculated for each trial. Additionally, the mean duration of detection events per trial was calculated to represent the average time available for a predator to respond to the prey.

### Statistical analysis

Each behavioural metric per trial, i.e. the proportion of frames that exceeded the threshold, mean magnitude within these frames, number of detection events and mean duration of detection events, was included as a response variable in a separate set of generalised linear mixed effect models (GLMMs) using the ‘glmmTMB’ R package v.1.1.7 (Brooks *et al*., 2017). The models for the proportion of frames were fitted with a beta distribution, the models for the mean magnitude in frames that exceeded the threshold were fitted using a gamma distribution with log link function, the models for the number of detection events were fitted with a negative binomial distribution, and the models for mean duration of detection events a Gaussian distribution. Model assumptions were checked for each distribution type: homogeneity of variance (gamma, Gaussian), dispersion of variance (beta, negative binomial), and normality of residuals (Gaussian) by visual inspection of diagnostic plots (i.e. QQ and dispersion plots) with R package ‘DHARMa’ v.0.4.6 (Hartig & Lohse, 2022).

A null model was built for each response variable including only the random effect of guppy tank of origin (i.e. their holding tank). The null model was then compared to seven alternative models each including different explanatory variables: temperature as the only main effect, turbidity as the only main effect, both temperature and turbidity as main effects (Temperature + Turbidity), an interaction term between temperature and turbidity as well as the individual main effects (Temperature × Turbidity), number of guppies included in each trial, trial order (1 to 4) to account for repeated testing (MacGregor & Ioannou, 2021), and minutes from midnight to account for diel variations in fish activity levels (Table 1 and 2). To improve interpretability and mitigate the risk of overfitting, we assessed the effects of control variables, i.e. the number of guppies, trial order, and minutes from midnight, through separate models. By opting for separate models, we aim to achieve clearer insights into the individual effects of control variables on the response variables, maintaining a balance between model simplicity and the need to understand the impact of various factors. The likelihood of each model was then compared using the difference in the Akaike Information Criterion corrected for small sample size (ΔAICc) from the best fitting model. Ranking models with different explanatory variables by their ΔAICc value allows for the inference of which variables drive variation in the response. If the difference in the ΔAICc is greater than 2 units, the model with the smaller ΔAICc value is considered to have strong support as being more likely given the data (Burnham & Anderson, 2002). When models fall within 0-2 units of the most likely model, it indicates a similar fit. However, if a model with more parameters is within this range of a smaller, more-likely model, it suggests that the similarity in ΔAICc values is primarily due to the addition of an extra parameter, rather than a genuine improvement in fit (Burnham and Anderson, 2002). When the null model has the smallest ΔAICc value, it suggests that none of the explanatory variables explain variation in the response. The analysis was repeated on the 5 datasets derived from the different thresholds (the 99^th^, 95^th^, 90^th^, 80^th^ and 70^th^ percentiles of the aggregated mean magnitude values).

**Table 1.** Model comparisons for the proportion of frames that exceeded the threshold and mean magnitude within these frames. Each row represents an individual model g in the explanatory variables: the plus sign indicates multiple variables are main effects, while × indicates an interaction term between temperature and turbidity is ed as well as their main effects. Each model also included the random effect of holding tank. ΔAICc: Difference in AIC value corrected for small sample size between model and the most likely model. df: number of components for each model. In red are highlighted the null models and in bold the models with greatest support dering both the ΔAICc and df).

| 99<sup>th</sup> | ΔAICc | df | 95<sup>th</sup> | ΔAICc | df | 90<sup>th</sup> | ΔAICc | df | 80<sup>th</sup> | ΔAICc | df | 70<sup>th</sup> | ΔAICc | df |
| --- | --- | --- | --- | --- | --- | --- | --- | --- | --- | --- | --- | --- | --- | --- |
| **Proportion of frames that exceeded the threshold** |  |  |  |  |  |  |  |  |  |  |  |  |  |  |
| **Repeated Testing** | **0.0** | **4** | **Turbidity** | **0.0** | **4** | Temp. × Turb. | 0.0 | 6 | Temp. × Turb. | 0.0 | 6 | **Temp. + Turb.** | **0.0** | **5** |
| Turbidity | 7.7 | 4 | Temp. × Turb. | 0.7 | 6 | **Temp. + Turb.** | **1.2** | **5** | **Temp. + Turb.** | **0.0** | **5** | Temp. × Turb. | 0.3 | 6 |
| Temp. + Turb. | 8.3 | 5 | Repeated Testing | 1.3 | 4 | Turbidity | 2.3 | 4 | Turbidity | 2.5 | 4 | Turbidity | 2.7 | 4 |
| Temp. × Turb. | 8.4 | 6 | Temp. + Turb. | 1.6 | 5 | Min from Midnight | 14.4 | 4 | Min from Midnight | 11.9 | 4 | Min from Midnight | 12.8 | 4 |
| Min from Midnight | 8.6 | 4 | Min from Midnight | 4.9 | 4 | Repeated Testing | 19.2 | 4 | Temperature | 15.5 | 4 | Temperature | 15.1 | 4 |
| *Null Model* | *13.3* | *3* | *Null Model* | *11.6* | *3* | *Null Model* | *20.4* | *3* | *Null Model* | *16.8* | *3* | *Null Model* | *16.9* | *3* |
| Temperature | 13.8 | 4 | Temperature | 13.4 | 4 | Temperature | 20.5 | 4 | Repeated Testing | 18.4 | 4 | Number of Fish | 18.9 | 4 |
| Number of Fish | 14.9 | 4 | Number of Fish | 13.7 | 4 | Number of Fish | 22.4 | 4 | Number of Fish | 18.8 | 4 | Repeated Testing | 19.1 | 4 |
| **Mean magnitude within frames that exceeded the threshold** |  |  |  |  |  |  |  |  |  |  |  |  |  |  |
| **Temperature** | **0.0** | **4** | **Temperature** | **0.0** | **4** | **Temp. + Turb.** | **0.0** | **5** | **Temperature** | **0.0** | **4** | **Repeated Testing** | **0.0** | **4** |
| Temp. + Turb. | 2.2 | 5 | Temp. + Turb. | 1.3 | 5 | Temp. × Turb. | 2.1 | 6 | **Repeated Testing** | **0.6** | **4** | Temperature | 4.6 | 4 |
| Temp. × Turb. | 4.5 | 6 | Temp. × Turb. | 3.5 | 6 | Temperature | 3.7 | 4 | Temp. + Turb. | 1.3 | 5 | Temp. × Turb. | 5.6 | 6 |
| *Null Model* | *17.1* | *3* | *Null Model* | *26.8* | *3* | Repeated Testing | 5.9 | 4 | Temp. × Turb. | 2.5 | 6 | Temp. + Turb. | 6.1 | 5 |
| Min from Midnight | 18.4 | 4 | Repeated Testing | 27.7 | 4 | Turbidity | 14.4 | 4 | *Null Model* | *7.5* | *3* | *Null Model* | *9.3* | *3* |
| Repeated Testing | 18.7 | 4 | Turbidity | 28.0 | 4 | *Null Model* | *17.7* | *3* | Turbidity | 8.6 | 4 | Turbidity | 10.7 | 4 |
| Number of Fish | 19.2 | 4 | Number of Fish | 28.3 | 4 | Min from Midnight | 19.9 | 4 | Min from Midnight | 9.6 | 4 | Min from Midnight | 11.2 | 4 |
| Turbidity | 19.3 | 4 | Min from Midnight | 29.0 | 4 | Number of Fish | 19.9 | 4 | Number of Fish | 9.6 | 4 | Number of Fish | 11.3 | 4 |

**Table 2.** Model comparisons for the number of detection events and mean duration of detection events. Each row represents an individual model varying in the atory variables: the plus sign indicates multiple variables are main effects, while × indicates an interaction term between temperature and turbidity is included as s their main effects. Each model also included the random effect of holding tank. ΔAICc: Difference in AIC between each model and the most likely model. df: number ponents for each model. In red are highlighted the null models and in bold the models with greatest support (considering both the ΔAICc and df).

| 99<sup>th</sup> | ΔAICc | df | 95<sup>th</sup> | ΔAICc | df | 90<sup>th</sup> | ΔAICc | df | 80<sup>th</sup> | ΔAICc | df | 70<sup>th</sup> | ΔAICc | df |
| --- | --- | --- | --- | --- | --- | --- | --- | --- | --- | --- | --- | --- | --- | --- |
| Number of detection events |  |  |  |  |  |  |  |  |  |  |  |  |  |  |
| Repeated Testing | 0.0 | 4 | Turbidity | 0.0 | 4 | Turbidity | 0.0 | 4 | Turbidity | 0.0 | 4 | Turbidity | 0.0 | 4 |
| Min from Midnight | 4.0 | 4 | Temp. + Turb. | 2.0 | 5 | Temp. + Turb. | 0.7 | 5 | Temp. + Turb. | 0.4 | 5 | Temp. + Turb. | 1.7 | 5 |
| Turbidity | 6.6 | 4 | Repeated Testing | 3.7 | 4 | Temp. × Turb. | 1.9 | 6 | Temp. × Turb. | 2.4 | 6 | Min from Midnight | 2.8 | 4 |
| Temp. + Turb. | 7.5 | 5 | Min from Midnight | 4.0 | 4 | Min from Midnight | 21.2 | 4 | Min from Midnight | 6.1 | 4 | Temp. × Turb. | 4.0 | 6 |
| Temp. × Turb. | 9.5 | 6 | Temp. × Turb. | 4.0 | 6 | Temperature | 28.2 | 4 | Temperature | 9.6 | 4 | *Null Model* | *4.3* | *3* |
| *Null Model* | *10.7* | *3* | *Null Model* | *12.9* | *3* | *Null Model* | *28.4* | *3* | *Null Model* | *9.6* | *3* | Repeated Testing | 5.3 | 4 |
| Temperature | 11.9 | 4 | Temperature | 14.6 | 4 | Repeated Testing | 30.1 | 4 | Repeated Testing | 10.8 | 4 | Temperature | 5.8 | 4 |
| Number of Fish | 12.9 | 4 | Number of Fish | 14.9 | 4 | Number of Fish | 30.6 | 4 | Number of Fish | 11.8 | 4 | Number of Fish | 6.4 | 4 |
| Mean duration of detection events |  |  |  |  |  |  |  |  |  |  |  |  |  |  |
| Temp. + Turb. | 0.0 | 5 | Min from Midnight | 0.0 | 4 | Repeated Testing | 0.0 | 4 | Repeated Testing | 0.0 | 4 | Temp. × Turb. | 0.0 | 6 |
| Number of Fish | 0.2 | 4 | Turbidity | 0.7 | 4 | Min from Midnight | 23.7 | 4 | Min from Midnight | 1.4 | 4 | Temp. + Turb. | 4.2 | 5 |
| Turbidity | 0.8 | 4 | Temp. + Turb. | 2.9 | 5 | *Null Model* | *26.9* | *3* | Turbidity | 2.1 | 4 | Turbidity | 4.8 | 4 |
| Temp. × Turb. | 2.3 | 6 | Temp. × Turb. | 3.5 | 6 | Temperature | 28.9 | 4 | Temp. × Turb. | 3.4 | 6 | Min from Midnight | 12.3 | 4 |
| Temperature | 2.7 | 4 | Number of Fish | 7.2 | 4 | Number of Fish | 29.0 | 4 | Temp. + Turb. | 4.2 | 5 | Temperature | 14.5 | 4 |
| *Null Model* | *3.2* | *3* | *Null Model* | *7.7* | *3* | Turbidity | 29.0 | 4 | *Null Model* | *5.0* | *3* | *Null Model* | *15.2* | *3* |
| Min from Midnight | 3.3 | 4 | Repeated Testing | 9.9 | 4 | Temp. + Turb. | 31.1 | 5 | Temperature | 7.0 | 4 | Repeated Testing | 17.4 | 4 |
| Repeated Testing | 4.2 | 4 | Temperature | 9.9 | 4 | Temp. × Turb. | 33.3 | 6 | Number of Fish | 7.2 | 4 | Number of Fish | 17.4 | 4 |

## Results

The proportion of frames per trial that exceeded the threshold was influenced mostly by turbidity, but also temperature at the lower thresholds. Models with only turbidity as the main effect (99^th^ and 95^th^ thresholds) and with temperature and turbidity as two main effects (90^th^, 80^th^ and 70^th^ thresholds) were much more likely than the null model (Table 1). For all thresholds, greater turbidity caused a reduction in the proportion of frames exceeding the threshold, suggesting that under these conditions guppies spent a greater time outside the field of view of the camera (Figure 4a).

**Figure 4.**
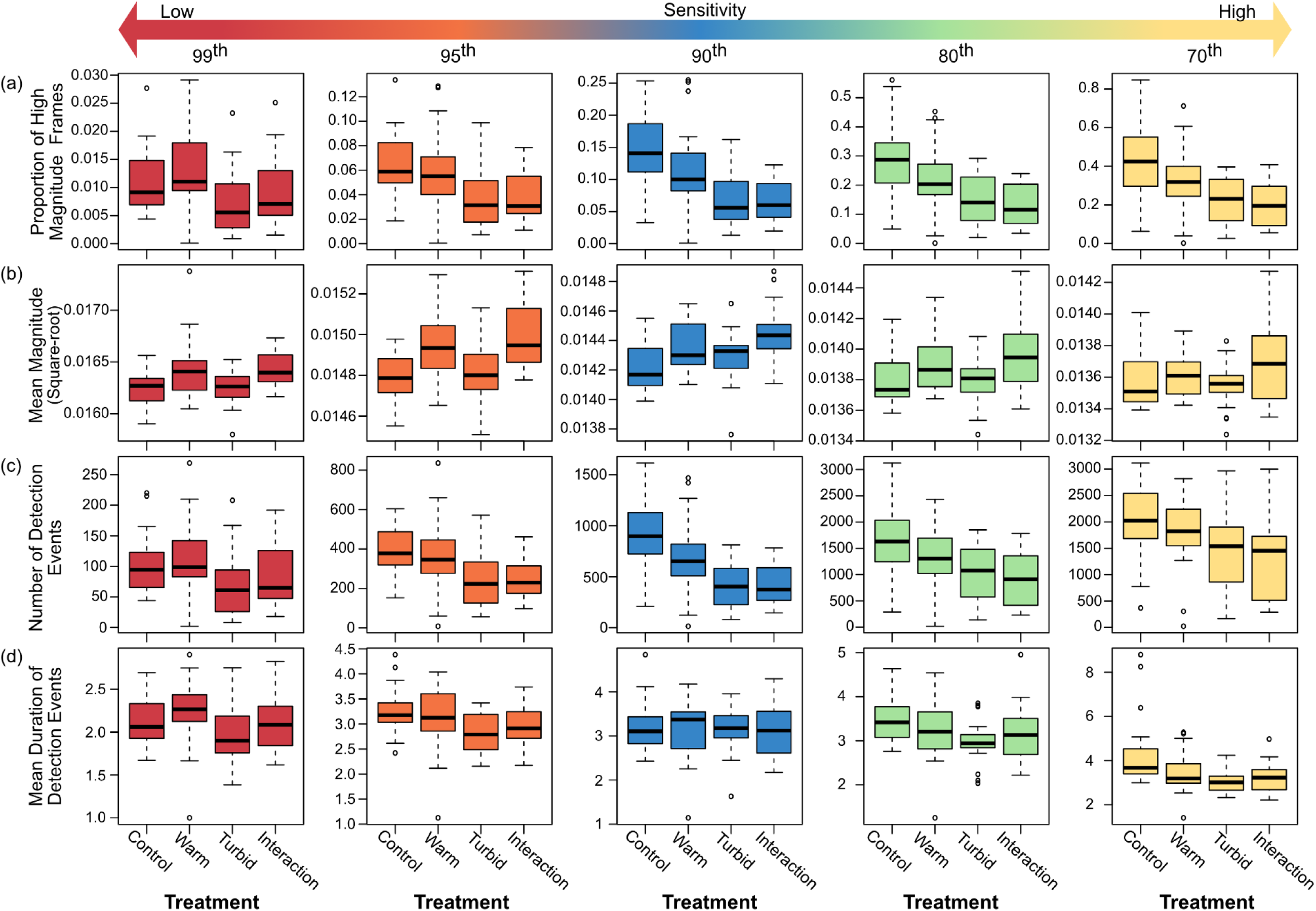
Response variables (**a**. Proportion of frames that exceeded the threshold, **b.** Mean magnitude within these frames, **c.** Number of detection events, **d.** Mean on of detection events) as a function of the four experimental treatments: control, warm, turbid and interaction, and derived from the different thresholds; note the nt scale of the y axes. In each plot, the horizontal black lines within the boxes represent the median value. The boxes span the interquartile range. The whiskers d from the most extreme data point by 1.5 times the interquartile range. The black circles represent outliers.

Temperature as a sole main effect was the leading predictor for the mean magnitude within frames that exceeded the threshold, as these models had a ΔAICc value of 0 for the 99^th^, 95^th^ and 80^th^ percentile thresholds datasets, and models with temperature were more likely than the null model. In the 90^th^ percentile threshold dataset, the model that included turbidity alongside temperature as additional main effect was the most likely model, and had considerably more support (ΔAICc = 3.7) than the model with temperature as the sole main effect (Table 1). In the 70^th^ percentile dataset, only the model with repeated testing as a main effect was more likely than the temperature-only model. Across thresholds, increases in temperature resulted in greater detectability (higher mean magnitudes) of fish within the view of the camera (Figure 4b).

The number of detection events was strongly reduced in turbid water. Specifically, the models that included turbidity as the sole main effect had a ΔAICc value of 0 for all thresholds except the 99^th^ percentile (Table 2); here the turbidity-only model was still more likely than the null model. In the interaction treatment, turbidity dominated the response with a similar number of detection events between the turbid and interaction treatments (Figure 4c).

Compared to the other response variables, there was greater variability between the different threshold datasets for the mean duration of detection events. The greater variability is due to higher thresholds dividing the longer detection events at low thresholds into multiple, shorter ones (Supplementary Figure 1d). The model including test order (i.e. repeated testing) as the main effect was the most likely for the 90^th^ and 80^th^ percentile thresholds, but it was less likely than the null model for the other thresholds. The model including the time of testing (i.e. minutes from midnight) was the most likely model for the 95^th^ percentile threshold, and was also supported (i.e. <2 ΔAICc units from the most likely model with same number of parameters) in the 80^th^ percentile threshold (Table 2). Models that included turbidity had support at all thresholds except the 90^th^ percentile, and in the 70^th^ percentile threshold dataset, there was strong support for the model with the temperature × turbidity interaction term. Increased turbidity reduced the mean duration of detection events (Figure 4d).

Test order and time of testing (i.e. repeated testing and minutes from midnight) had a varying degree of influence on the four behavioural responses and for the different threshold datasets (Table 1 and 2). In general, behavioural responses tended to increase with repeated testing (i.e. greater proportion of frames exceeding the threshold, higher mean magnitude within these frames, and more numerous and longer detection events: Supplementary Figure 2a and b), suggesting some level of habituation to the testing conditions where guppies became more active and approached the video camera more with repeated exposures to the experimental conditions. On the contrary, we observed a decrease in behaviours later in the day, possibly due to a decrease in overall activity with time of day (Supplementary Figure 2c). The number of fish swimming in the experimental arena for each trial did not influence any response, except the mean duration of detection events for the 99^th^ percentile threshold, where more guppies in the arena resulted in longer durations (Supplementary Figure 2c).

Lowering the threshold from the 99^th^ to the 70^th^ percentile increased the number of detection events almost 18-fold, from 8,461 to 149,767. This is consistent with the increase in the number of frames exceeding the threshold, which rose from 18,215 to 540,578. However, the most important predictors for the four response variables were relatively constant across the thresholds used (Table 1 and 2), supporting the consistency of the findings.

## Discussion

Our findings are consistent with previous research (Chaparro-Herrera et al., 2020; Jönsson et al., 2013; Turesson & Brönmark, 2007) showing that due to the visual impairment caused by turbid water, the proportion of time that prey were in close enough proximity to be detected by a predator decreased. Our study also demonstrates that once prey are close enough, recognising, targeting and capturing prey may become more difficult due to detection events being shorter in duration. However, we show that when detection events occurred in warmer water, prey detectability (i.e. the magnitude of the optical flow) was greater compared to the control and turbid treatments, likely due to the increased activity of the prey when within the camera’s field of view (Krause & Godin, 1995). These results suggest that in aquatic habitats that are becoming warmer and more turbid, warmer temperatures may, at least partially, compensate for the negative effects of turbidity on visual predators.

Predation risk can increase with warmer temperatures as ectothermic predators experience enhanced kinematics and greater predation drive (i.e. hunger levels caused by increased metabolic rates, Domenici et al., 2019). In response to increased threat, prey often reduce their activity levels to reduce their detectability (Ajemian et al., 2015; Eilam, 2005). However, prey fish are also subject to higher metabolic rates in warmer water (Bartolini et al., 2015). As a result, increased activity levels have been observed in prey fish under warmer conditions, even in the presence of a predator. Guppies show increased shoaling behaviour as a mitigating anti-predator strategy (Weetman et al., 1999). Although more aggregated prey can reduce encounter rates (Ioannou et al., 2011; Johannesen et al., 2014), our experiment did not find evidence supporting a change in the number or duration of detection events driven by temperature. A parallel study by Allibhai et al. (2023), which assessed the social behaviour of the guppies in this experiment using video recordings from above the experimental tank, reported an increase in shoaling tendencies at higher temperatures, but only at a local scale. Guppies were closer to their nearest neighbours in warmer water; however, the overall distances between all guppies in the arena were not affected by temperature. This suggests that although the guppies within each shoal were in closer proximity to one another at higher temperature, the distance between the shoals (which is more relevant to the frequency of encountering a predator) were not affected by temperature.

Our findings suggest that the frequency that prey were visible to the camera was dominated by the effects of turbidity, rather than by temperature. This is a comparative effect, i.e. where the response was dominated by only one of the multiple stressors (Folt et al., 1999). Previous research on guppy activity levels under turbid conditions has yielded mixed results, with some studies indicating decreased activity due to a heightened risk perception (Borner et al., 2015), while other studies show that turbidity has no influence on guppy activity levels (Zanghi et al., 2023), although this may be due to different levels of turbidity used in these experiments (e.g. 15 versus 700 NTU). Furthermore, observations by Allibhai et al. (2023) indicated that the increase in turbidity likely did not affect the risk perception of the guppies in this experiment, as refuge use remained unchanged. Nevertheless, the guppies exhibited reduced aggregation in turbid conditions, possibly due to the visual constraints of detecting distant shoal mates (MacGregor & Ioannou, 2023). While this reduction in aggregation could increase encounter rates (Ioannou et al., 2011; Johannesen et al., 2014), our findings suggest that this effect was outweighed by the direct effect of turbidity reducing the viewing distance from our hypothetical predator. In this context, a prey population that is more evenly distributed may have individuals closer to a predator compared to aggregated prey, yet remain undetected.

Quantifying the detectability of moving prey is more challenging compared to studying static prey, which are typically used in research on animal coloration (Stevens et al., 2007; Troscianko & Osorio, 2023). This complexity arises from the dynamic nature of motion, involving varying speed and changes in direction, which can influence detectability by predators (Cuthill et al., 2019). In our study, the overall number of detection events was greatly reduced (∼18 fold) with lower visual sensitivity (i.e. higher threshold of magnitude). This highlights the importance of considering different species’ visual abilities in a multi-predator system. However, modelling non-human visual abilities (such as spatial acuity, contrast sensitivity, and movement sensitivity) is also challenging (Caves & Johnsen, 2018). Our method of using optical flow to measure detectability offers several advantages, as optical flow algorithms capture the motion dynamics of animals, providing a continuous and objective measure of movement intensity. This approach can be valuable for predicting predator-prey interactions in more realistic settings. While there is still much to learn about the specific visual sensitivities of different predators (Heathcote et al., 2020), our method can incorporate species-specific visual acuity data to be tailored to specific systems.

As water temperature and turbidity increase in aquatic habitats globally due to human activities (e.g. quarrying (Ehlman et al., 2019), tilling (Dieter, 1991), agricultural eutrophication (Egertson et al., 2004), extreme weather events (Chiang et al., 2021; Mori et al., 2019), and global warming (Valipour et al., 2021)), there is a strong need to integrate behavioural studies with scenarios that better represent realistic natural conditions. The inclusion of multiple stressors in behavioural ecology has been increasing in current research. It is estimated that up to one-quarter of studies find interactions between stressors (i.e. synergies and antagonisms), in contrast to additive effects (Darling & Côté, 2008). Antagonistic interactions are more common than synergies, particularly in freshwater habitats (Darling et al., 2010; Jackson et al., 2016). While antagonistic interactions can be beneficial to impacted ecosystems due to mitigating effects, the overall impact can still be negative compared to pristine conditions (Orr et al., 2020; Vinebrooke et al., 2004). Our study did not observe interactions between stressors impacting the same response variable (e.g. duration of detection events). However, we found mitigating effects of increased temperature and turbidity on different metrics (i.e. mean magnitude and number of detection events) that together determine the detectability of prey.

Understanding shifts in animal behaviour as a result of environmental change can be a valuable monitoring tool (Bro-Jørgensen et al., 2019). Behavioural changes at an individual level can serve as early indicators of ecological stress, often leading to altered population dynamics and reproductive success (Candolin, 2009). Therefore, comprehensive and ecologically realistic approaches are important to inform the effective management and conservation of local natural resources.

## Ethical statement

All methods and procedures were performed in accordance with ASAB/ABS guidelines for the treatment of animals in behavioural research. The research was approved by the University of Bristol Animal Welfare and Ethics Review Body (UIN/21/003).

## Funding

This project was funded by the GW4 FRESH Centre for Doctoral Training in Freshwater Biosciences and Sustainability through an award to CZ (NE/R011524/1).

## Supplementary Figures

**Supplementary Figure 1.**
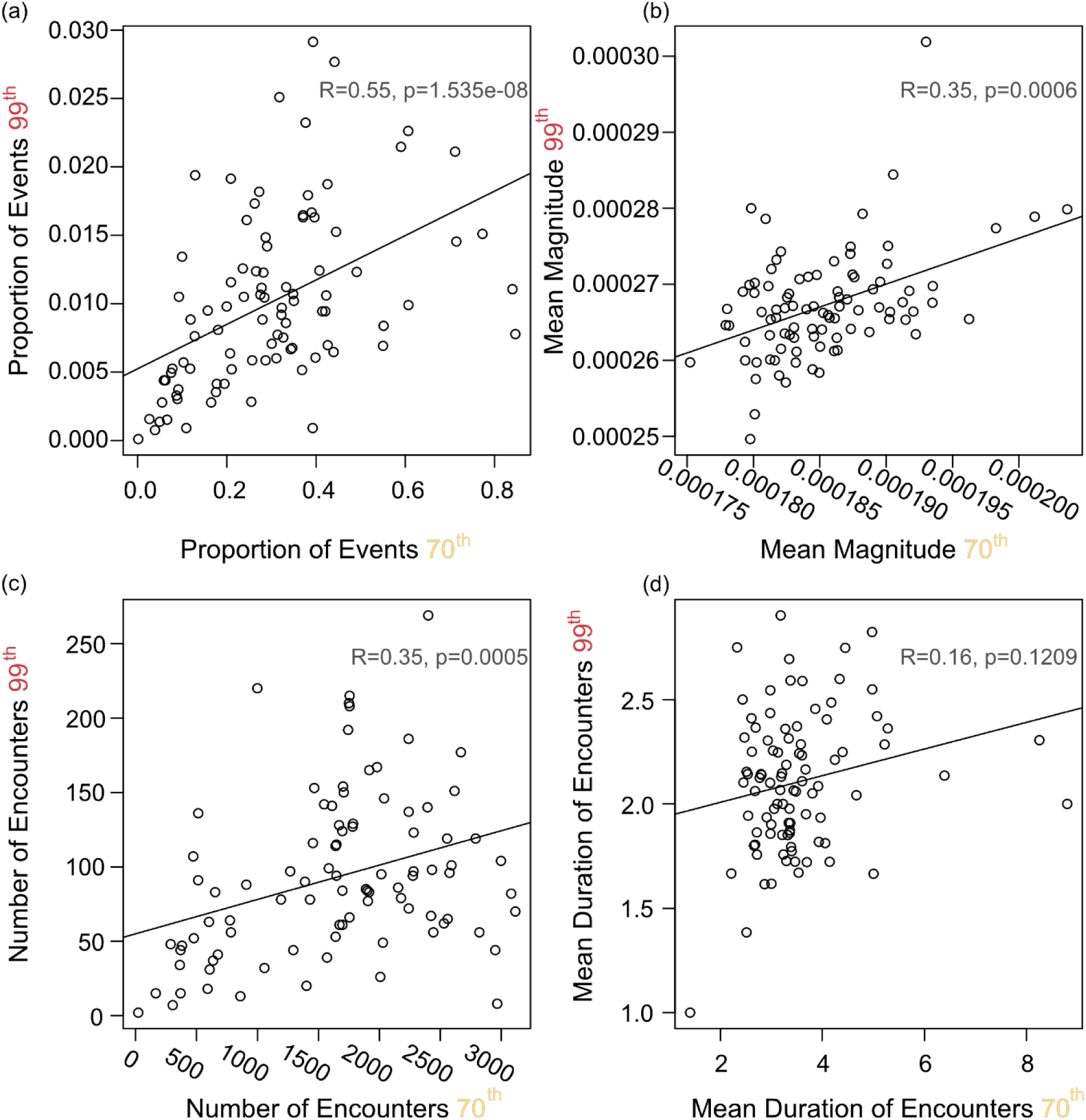
Relationship between the highest (99th percentile) and lowest (70^th^ percentile) sensitivity thresholds for each response variable: (a) proportion of frames that exceeded the threshold, (b) mean magnitude within these frames, (c) number of detection events, (d) mean duration of detection events (N=91). Regression lines, Spearman’s rank correlation coefficient (R) and significance of relationship (p) are reported for each response.

**Supplementary Figure 2.**
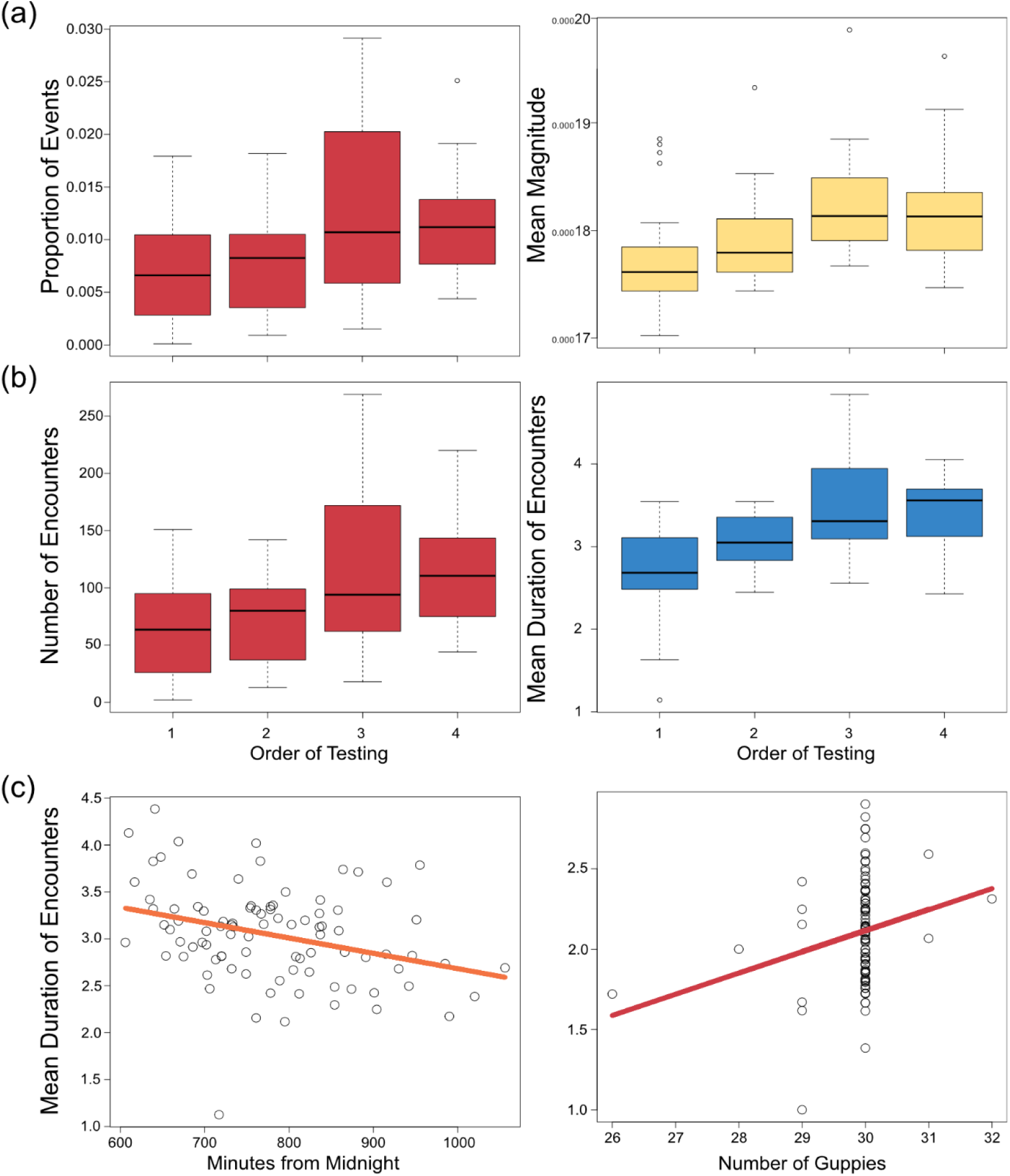
Response variables (a. left: proportion of frames that exceeded the threshold, a. right: mean magnitude within these frames, b. left: number of detection events, b. right: mean duration of detection events) as a function of the order of repeated testing (1 to 4), for the 99th (a. & b. left), 70th (a. right) and 80th (b. right) percentiles dataset. In each plot the horizontal black lines within the boxes represent the median value. The boxes span the interquartile range. The whiskers extend from the most extreme data point by 1.5 times the interquartile range. The black circles represent outliers. The mean duration of encounters as a function of (c) minutes from midnight (left, 95th percentile threshold) and number of guppies used in the trial (right, 99th percentile threshold). For each plot, dots represent individual observations (N=91), the lines represent the predicted values calculated from the GLMM coefficients. Note the different scale of the y axes across the different thresholds.

## Notes

### Competing Interest Statement

The authors have declared no competing interest.

## References

1. Abrahams, M. V., Mangel, M., & Hedges, K. (2007). Predator–prey interactions and changing environments: Who benefits? Philosophical Transactions of the Royal Society B: Biological Sciences, 362(1487), 2095–2104. 10.1098/rstb.2007.2102

2. Ajemian, M. J., Sohel, S., & Mattila, J. (2015). Effects of turbidity and habitat complexity on antipredator behavior of three-spined sticklebacks (Gasterosteus aculeatus): Antipredator behavior in sticklebacks. Environmental Biology of Fishes, 98(1), 45–55. 10.1007/s10641-014-0235-x

3. Alexander, E., Cai, L. T., Fuchs, S., Hladnik, T. C., Zhang, Y., Subramanian, V., Guilbeault, N. C., Vijayakumar, C., Arunachalam, M., Juntti, S. A., Thiele, T. R., Arrenberg, A. B., & Cooper, E. A. (2022). Optic flow in the natural habitats of zebrafish supports spatial biases in visual self-motion estimation. Current Biology, 32(23), 5008–5021.e8. 10.1016/j.cub.2022.10.009

4. Allibhai, I., Zanghi, C., How, M. J., & Ioannou, C. C. (2023). Increased water temperature and turbidity act independently to alter social behavior in guppies (Poecilia reticulata). Ecology and Evolution, 13(3), e9958. 10.1002/ece3.9958

5. Attwell, J. R., Ioannou, C. C., Reid, C. R., & Herbert-Read, J. E. (2021). Fish Avoid Visually Noisy Environments Where Prey Targeting Is Reduced. The American Naturalist, 198(3), 421–432. 10.1086/715434

6. Barron, J. L., Fleet, D. J., Beauchemin, S. S., & Burkitt, T. A. (1992). Performance of optical flow techniques. Proceedings 1992 IEEE Computer Society Conference on Computer Vision and Pattern Recognition, 236–242. 10.1109/CVPR.1992.223269

7. Bartolini, T., Butail, S., & Porfiri, M. (2015). Temperature influences sociality and activity of freshwater fish. Environmental Biology of Fishes, 98(3), 825–832. 10.1007/s10641-014-0318-8

8. Beauchamp, D. A., Baldwin, C. M., Vogel, J. L., & Gubala, C. P. (1999). Estimating diel, depth-specific foraging opportunities with a visual encounter rate model for pelagic piscivores. Canadian Journal of Fisheries and Aquatic Sciences, 56(S1), 128–139. 10.1139/f99-217

9. Borner, K. K., Krause, S., Mehner, T., Uusi-Heikkilä, S., Ramnarine, I. W., & Krause, J. (2015). Turbidity affects social dynamics in Trinidadian guppies. Behavioral Ecology and Sociobiology, 69(4), 645–651. 10.1007/s00265-015-1875-3

10. Bro-Jørgensen, J., Franks, D. W., & Meise, K. (2019). Linking behaviour to dynamics of populations and communities: Application of novel approaches in behavioural ecology to conservation. Philosophical Transactions of the Royal Society B: Biological Sciences, 374(1781), 20190008. 10.1098/rstb.2019.0008

11. Brooks, M. E., Kristensen, K., Benthem, K. J., van Magnusson, A., Berg, C. W., Nielsen, A., Skaug, H. J., Mächler, M., & Bolker, B. M. (2017). glmmTMB Balances Speed and Flexibility Among Packages for Zero-inflated Generalized Linear Mixed Modeling. The R Journal, 9(2), 378. 10.32614/RJ-2017-066

12. Bunkley, J. P., McClure, C. J. W., Kleist, N. J., Francis, C. D., & Barber, J. R. (2015). Anthropogenic noise alters bat activity levels and echolocation calls. Global Ecology and Conservation, 3, 62–71. 10.1016/j.gecco.2014.11.002

13. Burnham, K. P., & Anderson, D. R. (Eds.). (2002). Information and Likelihood Theory: A Basis for Model Selection and Inference. In Model Selection and Multimodel Inference: A Practical Information-Theoretic Approach (pp. 49–97). Springer. 10.1007/978-0-387-22456-5_2

14. Candolin, U. (2009). Population responses to anthropogenic disturbance: Lessons from three-spined sticklebacks Gasterosteus aculeatus in eutrophic habitats. Journal of Fish Biology, 75(8), 2108–2121. 10.1111/j.1095-8649.2009.02405.x

15. Carroll, J., & Sherratt, T. N. (2013). A direct comparison of the effectiveness of two anti-predator strategies under field conditions. Journal of Zoology, 291(4), 279–285. 10.1111/jzo.12074

16. Caves, E. M., & Johnsen, S. (2018). AcuityView: An r package for portraying the effects of visual acuity on scenes observed by an animal. Methods in Ecology and Evolution, 9(3), 793–797. 10.1111/2041-210X.12911

17. Chan, A. A. Y.-H., Giraldo-Perez, P., Smith, S., & Blumstein, D. T. (2010). Anthropogenic noise affects risk assessment and attention: The distracted prey hypothesis. Biology Letters, 6(4), 458–461. 10.1098/rsbl.2009.1081

18. Chaparro-Herrera, D. J., Nandini, S., & Sarma, S. S. S. (2020). Turbidity effects on feeding by larvae of the endemic Ambystoma mexicanum and the introduced Oreochromis niloticus in Lake Xochimilco, Mexico. Ecohydrology & Hydrobiology, 20(1), 91–101. 10.1016/j.ecohyd.2019.07.002

19. Chase, J. M., Abrams, P. A., Grover, J. P., Diehl, S., Chesson, P., Holt, R. D., Richards, S. A., Nisbet, R. M., & Case, T. J. (2002). The interaction between predation and competition: A review and synthesis. Ecology Letters, 5(2), 302–315. 10.1046/j.1461-0248.2002.00315.x

20. Chiang, F., Mazdiyasni, O., & AghaKouchak, A. (2021). Evidence of anthropogenic impacts on global drought frequency, duration, and intensity. Nature Communications, 12(1), Article 1. 10.1038/s41467-021-22314-w

21. Cuthill, I. C., Matchette, S. R., & Scott-Samuel, N. E. (2019). Camouflage in a dynamic world. Current Opinion in Behavioral Sciences, 30, 109–115. 10.1016/j.cobeha.2019.07.007

22. Darby, J., Clairbaux, M., Bennison, A., Quinn, J. L., & Jessopp, M. J. (2022). Underwater visibility constrains the foraging behaviour of a diving pelagic seabird. Proceedings of the Royal Society B: Biological Sciences, 289(1978), 20220862. 10.1098/rspb.2022.0862

23. Darling, E. S., & Côté, I. M. (2008). Quantifying the evidence for ecological synergies. Ecology Letters, 11(12), 1278–1286. 10.1111/j.1461-0248.2008.01243.x

24. Darling, E. S., McClanahan, T. R., & Côté, I. M. (2010). Combined effects of two stressors on Kenyan coral reefs are additive or antagonistic, not synergistic. Conservation Letters, 3(2), 122–130. 10.1111/j.1755-263X.2009.00089.x

25. Dawkins, M. S., Cain, R., & Roberts, S. J. (2012). Optical flow, flock behaviour and chicken welfare. Animal Behaviour, 84(1), 219–223. 10.1016/j.anbehav.2012.04.036

26. Dawkins, M. S., Lee, H., Waitt, C. D., & Roberts, S. J. (2009). Optical flow patterns in broiler chicken flocks as automated measures of behaviour and gait. Applied Animal Behaviour Science, 119(3), 203–209. 10.1016/j.applanim.2009.04.009

27. Dieter, C. D. (1991). Water Turbidity in Tilled and Untilled Prairie Wetlands. Journal of Freshwater Ecology, 6(2), 185–189. 10.1080/02705060.1991.9665292

28. Domenici, P., Allan, B. J. M., Lefrançois, C., & McCormick, M. I. (2019). The effect of climate change on the escape kinematics and performance of fishes: Implications for future predator–prey interactions. Conservation Physiology, 7(1), coz078. 10.1093/conphys/coz078

29. Domenici, P., Claireaux, G., & McKenzie, D. J. (2007). Environmental constraints upon locomotion and predator–prey interactions in aquatic organisms: An introduction. Philosophical Transactions of the Royal Society B: Biological Sciences, 362(1487), 1929–1936. 10.1098/rstb.2007.2078

30. Egertson, C. J., Kopaska, J. A., & Downing, J. A. (2004). A Century of Change in Macrophyte Abundance and Composition in Response to Agricultural Eutrophication. Hydrobiologia, 524(1), 145–156. 10.1023/B:HYDR.0000036129.40386.ce

31. Ehlman, S. M., Sandkam, B. A., Breden, F., & Sih, A. (2015). Developmental plasticity in vision and behavior may help guppies overcome increased turbidity. Journal of Comparative Physiology A, 201(12), 1125–1135. 10.1007/s00359-015-1041-4

32. Ehlman, S. M., Torresdal, J. D., & Fraser, D. F. (2019). Altered visual environment affects a tropical freshwater fish assemblage through impacts on predator–prey interactions. Freshwater Biology, 65(2), 316–324. 10.1111/fwb.13425

33. Eilam, D. (2005). Die hard: A blend of freezing and fleeing as a dynamic defense—implications for the control of defensive behavior. Neuroscience & Biobehavioral Reviews, 29(8), 1181–1191. 10.1016/j.neubiorev.2005.03.027

34. Einfalt, L. M., & Wahl, D. H. (1997). Prey selection by juvenile walleye as influenced by prey morphology and behavior. Canadian Journal of Fisheries and Aquatic Sciences, 54(11), 2618– 2626. 10.1139/f97-172

35. Ferrero, D. M., Lemon, J. K., Fluegge, D., Pashkovski, S. L., Korzan, W. J., Datta, S. R., Spehr, M., Fendt, M., & Liberles, S. D. (2011). Detection and avoidance of a carnivore odor by prey. Proceedings of the National Academy of Sciences, 108(27), 11235–11240. 10.1073/pnas.1103317108

36. Folt, C. L., Chen, C. Y., Moore, M. V., & Burnaford, J. (1999). Synergism and antagonism among multiple stressors. Limnology and Oceanography, 44(3part2), 864–877. 10.4319/lo.1999.44.3_part_2.0864

37. Franco, M. F., Santostefano, F., Kelly, C. D., & Montiglio, P.-O. (2022). Studying predator foraging mode and hunting success at the individual level with an online videogame. Behavioral Ecology, 33(5), 967–978. 10.1093/beheco/arac063

38. Gregory, R. S. (1993). Effect of Turbidity on the Predator Avoidance Behaviour of Juvenile Chinook Salmon (*Oncorhynchus tshawytscha*). Canadian Journal of Fisheries and Aquatic Sciences, 50(2), 241–246. 10.1139/f93-027

39. Hansen, A. G., Beauchamp, D. A., & Schoen, E. R. (2013). Visual Prey Detection Responses of Piscivorous Trout and Salmon: Effects of Light,Turbidity,and Prey Size. Transactions of the American Fisheries Society, 142(3), 854–867. 10.1080/00028487.2013.785978

40. Harris, S., Ramnarine, I. W., Smith, H. G., & Pettersson, L. B. (2010). Picking personalities apart: Estimating the influence of predation, sex and body size on boldness in the guppy Poecilia reticulata. Oikos, 119(11), 1711–1718. 10.1111/j.1600-0706.2010.18028.x

41. Hartig, F., & Lohse, L. (2022). *DHARMa: Residual Diagnostics for Hierarchical (Multi-Level / Mixed) Regression Models* (Version 0.4.6) [R]. https://CRAN.R-project.org/package=DHARMa

42. Heathcote, R. J. P., Troscianko, J., Darden, S. K., Naisbett-Jones, L. C., Laker, P. R., Brown, A. M., Ramnarine, I. W., Walker, J., & Croft, D. P. (2020). A Matador-like Predator Diversion Strategy Driven by Conspicuous Coloration in Guppies. Current Biology, 30(14), 2844–2851.e8. 10.1016/j.cub.2020.05.017

43. Ioannou, C. C., Bartumeus, F., Krause, J., & Ruxton, G. D. (2011). Unified effects of aggregation reveal larger prey groups take longer to find. Proceedings of the Royal Society B: Biological Sciences, 278(1720), 2985–2990. 10.1098/rspb.2011.0003

44. Ioannou, C. C., & Krause, J. (2009). Interactions between background matching and motion during visual detection can explain why cryptic animals keep still. Biology Letters, 5(2), 191–193. 10.1098/rsbl.2008.0758

45. Ioannou, C. C., Ruxton, G., & Krause, J. (2008). Search rate, attack probability, and the relationship between prey density and prey encounter rate. Behavioral Ecology, 19, 842–846. 10.1093/beheco/arn038

46. Jackson, M. C., Loewen, C. J. G., Vinebrooke, R. D., & Chimimba, C. T. (2016). Net effects of multiple stressors in freshwater ecosystems: A meta-analysis. Global Change Biology, 22(1), 180–189. 10.1111/gcb.13028

47. Johannesen, A., Dunn, A. M., & Morrell, L. J. (2014). Prey aggregation is an effective olfactory predator avoidance strategy. PeerJ, 2, e408. 10.7717/peerj.408

48. Jönsson, M., Ranåker, L., Nilsson, P. A., & Brönmark, C. (2013). Foraging efficiency and prey selectivity in a visual predator: Differential effects of turbid and humic water. Canadian Journal of Fisheries and Aquatic Sciences, 70(12), 1685–1690. 10.1139/cjfas-2013-0150

49. Khorrami, P., Wang, J., & Huang, T. (2012). Multiple Animal Species Detection Using Robust Principal Component Analysis and Large Displacement Optical Flow. ICPR Workshop on Visual Observation andAnalysis of Animal and Insect Behavior (VAIB).

50. Krause, J., & Godin, J.-G. (1996). Influence of prey foraging posture on flight behavior and predation risk: Predators take advantage of unwary prey. Behavioral Ecology - BEHAV ECOL, 7, 264–271. 10.1093/beheco/7.3.264

51. Krause, J., & Godin, J.-G. J. (1995). Predator preferences for attacking particular prey group sizes: Consequences for predator hunting success and prey predation risk. Animal Behaviour, 50(2), 465–473. 10.1006/anbe.1995.0260

52. Lima, S. L. (1998). Nonlethal Effects in the Ecology of Predator-Prey Interactions. BioScience, 48(1), 25–34. 10.2307/1313225

53. Lima, S. L., & Dill, L. M. (1990). Behavioral decisions made under the risk of predation: A review and prospectus. Canadian Journal of Zoology, 68(4), 619–640. 10.1139/z90-092

54. MacGregor, H. E. A., & Ioannou, C. C. (2021). Collective motion diminishes, but variation between groups emerges, through time in fish shoals. Royal Society Open Science, 8(10), 210655. 10.1098/rsos.210655

55. MacGregor, H. E. A., & Ioannou, C. C. (2023). Shoaling behaviour in response to turbidity in three-spined sticklebacks. Ecology and Evolution, 13(11), e10708. 10.1002/ece3.10708

56. Magurran, A. E. (1990). The adaptive significance of schooling as an anti-predator defence in fish. Ann. Zool. Fennici., 27, 51–66.

57. Mori, N., Shimura, T., Yoshida, K., Mizuta, R., Okada, Y., Fujita, M., Khujanazarov, T., & Nakakita, E. (2019). Future changes in extreme storm surges based on mega-ensemble projection using 60-km resolution atmospheric global circulation model. Coastal Engineering Journal, 61(3), 295–307. 10.1080/21664250.2019.1586290

58. Morris-Drake, A., Kern, J. M., & Radford, A. N. (2016). Cross-modal impacts of anthropogenic noise on information use. Current Biology, 26(20), R911–R912. 10.1016/j.cub.2016.08.064

59. Nieman, C. L., & Gray, S. M. (2019). Visual performance impaired by elevated sedimentary and algal turbidity in walleye *Sander vitreus* and emerald shiner *Notropis atherinoides*. Journal of Fish Biology, 95(1), 186–199. 10.1111/jfb.13878

60. Orr, J. A., Vinebrooke, R. D., Jackson, M. C., Kroeker, K. J., Kordas, R. L., Mantyka-Pringle, C., Van den Brink, P. J., De Laender, F., Stoks, R., Holmstrup, M., Matthaei, C. D., Monk, W. A., Penk, M. R., Leuzinger, S., Schäfer, R. B., & Piggott, J. J. (2020). Towards a unified study of multiple stressors: Divisions and common goals across research disciplines. Proceedings of the Royal Society B: Biological Sciences, 287(1926), 20200421. 10.1098/rspb.2020.0421

61. Paine, R. T., Tegner, M. J., & Johnson, E. A. (1998). Compounded Perturbations Yield Ecological Surprises. Ecosystems, 1(6), 535–545. 10.1007/s100219900049

62. Ruxton, G. D., Allen, W. L., Sherratt, T. N., & Speed, M. P. (2018). Batesian mimicry and masquerade. In G. D. Ruxton, W. L. Allen, T. N. Sherratt, & M. P. Speed (Eds.), Avoiding Attack: The Evolutionary Ecology of Crypsis, Aposematism, and Mimicry (p. 0). Oxford University Press. 10.1093/oso/9780199688678.003.0010

63. Sims, D. W., Southall, E. J., Humphries, N. E., Hays, G. C., Bradshaw, C. J. A., Pitchford, J. W., James, A., Ahmed, M. Z., Brierley, A. S., Hindell, M. A., Morritt, D., Musyl, M. K., Righton, D., Shepard, E. L. C., Wearmouth, V. J., Wilson, R. P., Witt, M. J., & Metcalfe, J. D. (2008). Scaling laws of marine predator search behaviour. Nature, 451(7182), Article 7182. 10.1038/nature06518

64. Stevens, M., Párraga, C. A., Cuthill, I. C., Partridge, J. C., & Troscianko, T. S. (2007). Using digital photography to study animal coloration: USING CAMERAS TO STUDY ANIMAL COLORATION. Biological Journal of the Linnean Society, 90(2), 211–237. 10.1111/j.1095-8312.2007.00725.x

65. Sun, D., Roth, S., & Black, M. J. (2014). A Quantitative Analysis of Current Practices in Optical Flow Estimation and the Principles Behind Them. International Journal of Computer Vision, 106(2), 115–137. 10.1007/s11263-013-0644-x

66. Szopa-Comley, A. W., Duffield, C., Ramnarine, I. W., & Ioannou, C. C. (2020). Predatory behaviour as a personality trait in a wild fish population. Animal Behaviour, 170, 51–64. 10.1016/j.anbehav.2020.10.002

67. The MathWorks Inc. (2021). *MATLAB R2021a* (Version 9.10.0.1739362 (R2021a) Update 5) [Computer software]. Natick, Massachusetts: The MathWorks Inc. https://www.mathworks.com

68. Townsend, C. R., Uhlmann, S. S., & Matthaei, C. D. (2008). Individual and combined responses of stream ecosystems to multiple stressors. Journal of Applied Ecology, 45(6), 1810–1819. 10.1111/j.1365-2664.2008.01548.x

69. Troscianko, J., & Osorio, D. (2023). A model of colour appearance based on efficient coding of natural images. PLOS Computational Biology, 19(6), e1011117. 10.1371/journal.pcbi.1011117

70. Troscianko, T., Benton, C. P., Lovell, P. G., Tolhurst, D. J., & Pizlo, Z. (2009). Camouflage and visual perception. 364, 449–461. 10.1098/rstb.2008.0218

71. Turesson, H., & Brönmark, C. (2007). Predator–prey encounter rates in freshwater piscivores: Effects of prey density and water transparency. Oecologia, 153(2), 281–290. 10.1007/s00442-007-0728-9

72. Turesson, H., Satta, A., & Domenici, P. (2009). Preparing for escape: Anti-predator posture and fast-start performance in gobies. Journal of Experimental Biology, 212(18), 2925–2933. 10.1242/jeb.032953

73. Utne-Palm, A. C. (2002). Visual feeding of fish in a turbid environment: Physical and behavioural aspects. Marine and Freshwater Behaviour and Physiology, 35(1–2), 111–128. 10.1080/10236240290025644

74. Valipour, M., Bateni, S. M., & Jun, C. (2021). Global Surface Temperature: A New Insight. Climate, 9(5), 81. 10.3390/cli9050081

75. Vinebrooke, R. D., Cottingham, K. L., Norberg, Marten Scheffer, J., Dodson, J. J., Maberly, S. C., & Sommer, U. (2004). Impacts of multiple stressors on biodiversity and ecosystem functioning: The role of species co-tolerance. Oikos, 104(3), 451–457. 10.1111/j.0030-1299.2004.13255.x

76. Vogel, J. L., & Beauchamp, D. A. (1999). Effects of Light Prey Size and Turbidity on Reaction distances of lake trout. Canadian Journal of Fisheries and Aquatic Sciences, 56, 1293–1297.

77. Wearmouth, V. J., McHugh, M. J., Humphries, N. E., Naegelen, A., Ahmed, M. Z., Southall, E. J., Reynolds, A. M., & Sims, D. W. (2014). Scaling laws of ambush predator ‘waiting’ behaviour are tuned to a common ecology. Proceedings of the Royal Society B: Biological Sciences, 9.

78. Weetman, D., Atkinson, D., & Chubb, J. C. (1998). Effects of temperature on anti-predator behaviour in the guppy,Poecilia reticulata. Animal Behaviour, 55(5), 1361–1372. 10.1006/anbe.1997.0666

79. Weetman, D., Atkinson, D., & Chubb, J. C. (1999). Water temperature influences the shoaling decisions of guppies, Poecilia reticulata, under predation threat. Animal Behaviour, 58(4), 735–741. 10.1006/anbe.1999.1191

80. Zanghi, C., Munro, M., & Ioannou, C. C. (2023). Temperature and turbidity interact synergistically to alter anti-predator behaviour in the Trinidadian guppy. Proceedings of the Royal Society B: Biological Sciences, 290(2002), 20230961. 10.1098/rspb.2023.0961

81. Zanghi, C., Penry-Williams, I. L., Genner, M. J., Deacon, A. E., & Ioannou, C. C. (2024). Multiple environmental stressors affect predation pressure in a tropical freshwater system. Communications Biology, 7(1), 1–11. 10.1038/s42003-024-06364-6

